# A non-invasive eDNA tool for detecting sea lamprey larvae in river sediments: analytical validation and field testing in a low abundance ecosystem

**DOI:** 10.1101/2020.02.08.938803

**Authors:** Miguel Baltazar-Soares, Adrian C. Pinder, Andrew J. Harrison, Will Oliver, Jessica Picken, J. Robert Britton, Demetra Andreou

## Abstract

Anthropogenic activities are increasingly threatening aquatic biodiversity, especially anadromous species. Monitoring and conservation measures are thus required to protect, maintain, and restore imperilled populations. While many species can be surveyed using traditional capture and visual census techniques, species that use riverine habitats in a less conspicuous manner, such as sea lamprey *Petromyzon marinus*, can be more challenging to monitor. Sea lamprey larvae (ammocoetes) can spend several years in freshwater burrowed within soft sediments, inhibiting their detection and assessment. Here we present an environmental DNA (eDNA) assay to detect ammocoetes burrowed in the sediment. We performed a battery of tests that ensured both species-specificity of the assay as well as the capacity to detect ammocoetes when abundances are low. Experiments on burrowing activity suggested that most of the DNA released into the sediment occurs during burrowing. Overall, we demonstrated this new molecular-based tool is an efficient and effective complement to traditional monitoring activities targeting larval stages of sea lampreys.

## Introduction

Anthropogenic activities in freshwater ecosystems can result in deleterious effects on biodiversity (Dodds, Perkin, & Gerken, 2013), including on those species with complex life cycles, such as anadromous fishes (Dias et al., 2017). Effective conservation management of freshwater biodiversity is reliant on robust monitoring that enables long-term spatial and temporal patterns to be detected (Radinger et al., 2019). This monitoring can, however, be challenging in many freshwater ecosystems, with biases in sampling methods leading to issues associated with false negative data, especially if relying on visual detection and counting (Thomsen et al., 2012). Threatened species that have complex life cycles, and with life-stages that occupy a range of different habitats, can then present further challenges to sampling efficacy within monitoring programmes (Radinger et al., 2019).

An alternative to the application of capture and visual sampling methods is the use of environmental DNA (eDNA) methods that are designed to detect the DNA of organisms within environmental samples, such as water (Thomsen & Willerslev, 2015a). Although eDNA based detection methods require extensive validation prior to their application in the field, their capacity to detect DNA from environmental samples without capturing or visually confirming the species presence is becoming increasingly cost-efficient (Doi et al., 2017; Evans, Shirey, Wieringa, Mahon, & Lamberti, 2017; Ficetola, Miaud, Pompanon, & Taberlet, 2008). Correspondingly, the application of eDNA detection to monitoring biodiversity is now well established, with a broad range of applications, including species detection in either an ancient or contemporary context, from the reconstruction of past faunal or floral assemblages to the identification of range expansions / biological invasions (Bohmann et al., 2014). In the freshwater environment, it has been used to document habitat utilisation, detect the presence of rare and invasive species, and quantify spawning activity in pre-defined areas (Bracken, Rooney, Kelly□Quinn, King, & Carlsson, 2019; Stoeckle, Soboleva, & Charlop-Powers, 2017; Thomsen & Willerslev, 2015b), including the extent of upstream spawning migrations in anadromous fish species (Antognazza et al., 2019).

The sea lamprey *Petromyzon marinus* is a jawless vertebrate, one of the few surviving species of this ancient group of animals (Guo, Andreou, & Britton, 2017). Its complex life cycle comprises stages in fresh, brackish and saltwater. Mating occurs in freshwater in primitive nests built by males to attract females (Guo et al., 2017). Batches of eggs are deposited within the gravel substrate where they reside for an extended larval phase (∼5-6 years) until they emerge during the ontogenetic metamorphosis from larvae (ammocoetes) to young juveniles (transformers) which then drift downstream to deeper areas of low water velocity, before settling into areas of soft sediment (Pinder, Hopkins, Scott, & Britton, 2016). Their downstream migration to sea is accomplished by successive settlement / emergence events; a process lasting at least five years (Quintella, Andrade, Espanhol, & Almeida, 2005). In the sea, adults are parasitic on fish until sexual maturity when they then return to freshwater (Sorensen & Vrieze, 2003).

In Europe threats to its conservation status include river fragmentation, habitat loss and declining water quality (Pinder et al., 2016), resulting in protected status through designation under Annex II of the EU Habitats Directive (Directive 92/43/EEC). Robust population monitoring is required to ensure that populations achieve the regulatory target of ‘favourable conservation status’, with monitoring strategies usually targeting the freshwater stages, especially nest counts during spawning activity or abundance estimates of ammocoetes in the sediment (Guo et al., 2017; Mateus, Rodríguez-Muñoz, Quintella, Alves, & Almeida, 2012). As ammocoetes remain in freshwater for several years, targeting sediment sites to infer ammocoete presence and / or abundance could provide evidence for the sustainability of sea lamprey populations within specific rivers and habitats. Correspondingly, the aim here was to develop a molecular-based assay designed and tested to capture DNA molecules of sea lamprey in sediment samples, in order to provide a conservation monitoring tool with the capability of being utilised all year round.

## Material and methods

### Sampling site and identification of sediment deposition sites

This work was performed on the River Frome, Southern England, where adults spawn in the lower reaches, but with considerable inter-annual variation in numbers (Pinder et al., 2016). Sea lamprey ammocoete distribution in the River Frome is largely unknown, therefore considerable effort was allocated to identifying suitable sedimentation sites. Focusing on river meanders - where the likelihood of soft sediment deposition is higher - five sites where ammocoetes have been found in previous years were chosen along a 13 km stretch of the lower River Frome (Fig.1a).

**Figure 1.**
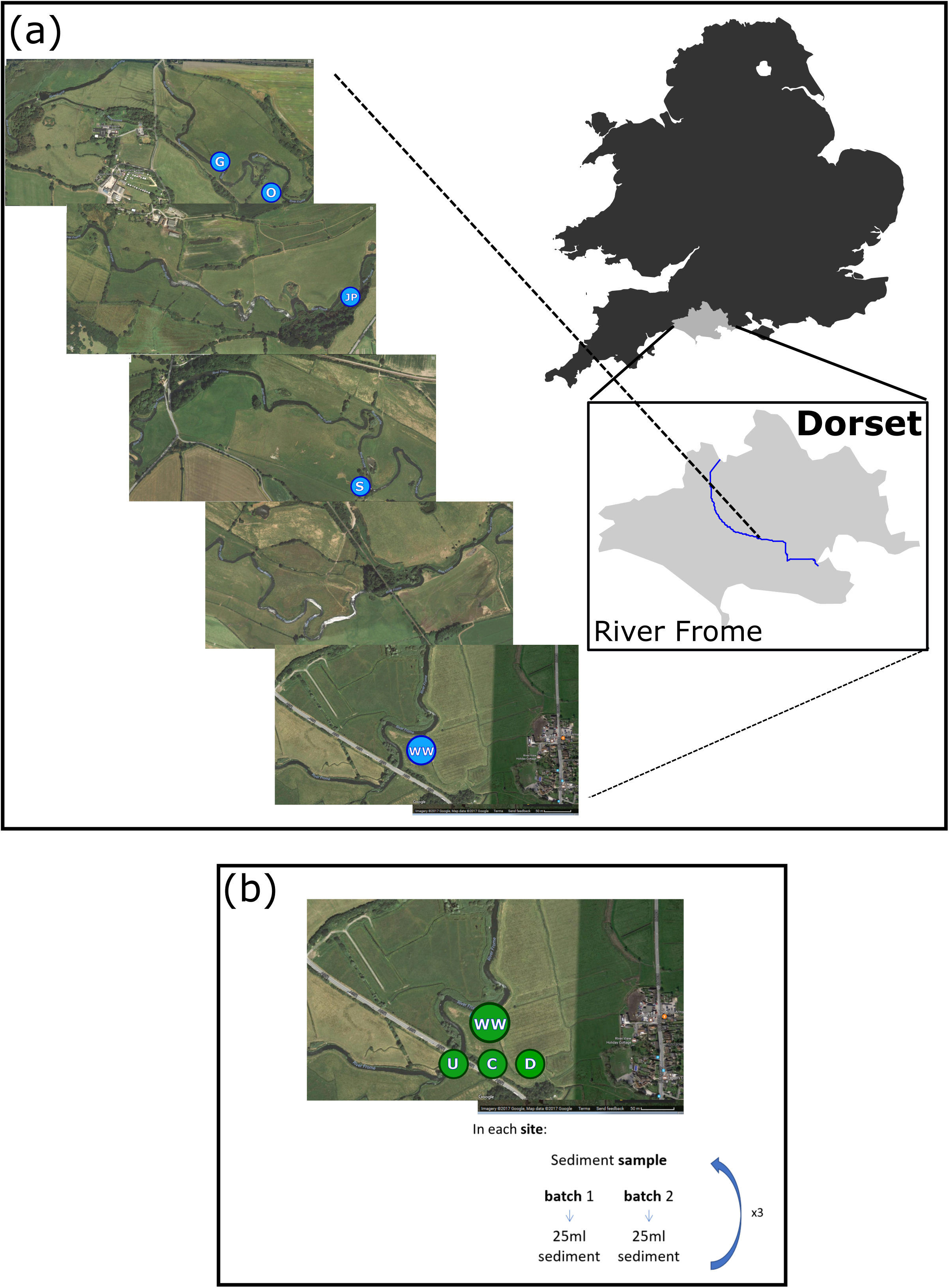
Sampling sites and strategy for environmental DNA screens. (a) Riverside sites of sediment sampling along a transect of the river Frome. (b) Sampling strategy adopted for the monitoring *in situ* strategy at the five sampling sites. Here, *U* stands for the location upstream of the site, *C* stands for the location at the site and *D* stands for the location downstream of the sites.

### Identification of sea lamprey ammocoetes via mitochondrial DNA barcoding

With the non-migratory brook lamprey *Lampetra planeri* being prevalent throughout the river system, a primary requirement was validation of the ability to distinguish between brook and sea lamprey. Ammocoetes of sea and brook lamprey are virtually indistinguishable prior to the onset of pigmentation but, thereafter, morphological discrimination is possible based on pigmentation around the oral hood and on the caudal fin, where it is present in sea lamprey but not in brook lamprey (Kelly & King, 2001).

For the identification of sea versus brook lamprey ammocoetes, we collected and euthanised six lamprey ammocoetes, three pigmented (putative sea lamprey) and three without pigment (putative brook lamprey). Species identification was verified with mitochondrial DNA (mtDNA) barcodes. For this, a fraction of the overall body tissue was used for the DNA extraction, amplification and sequencing. Extraction of DNA was performed with a commercially available extraction kit (Qiagen^©^) and followed the manufacturer protocol. Amplification targeted a portion of the control region in the mitochondrial DNA, using a mix of primers from two different studies that were specifically designed for brook and sea lamprey (Almada et al., 2008; Rodríguez□Muñoz, Waldman, Grunwald, Roy, & Wirgin, 2004). Amplification was through Polymerase Chain Reaction (PCR) under the following conditions: 95°C for 15 minutes for *Taq* polymerase activation, followed by 35 cycles of a denaturation step at 94 °C for 30 s, annealing step at 55 °C for 30 s and elongation step at 72 °C for 30 s, and a final elongation step at 72 °C for 10 minutes. The PCR mix was composed of 5 ul of QIAGEN MasterMix, 0.4 ul of forward and reverse primer (5 uM), 2.6 ul purified water and 2 ul DNA (∼100 ng·ul^-1^). Successful amplification was verified in agarose gels (1%) submitted to 70 V for 45 minutes. Sequencing was outsourced to GENEWIZ^©^; sequences were aligned and curated manually in BioEdit (Hall, 1999) and compared against those deposited in the National Center for Biotechnology Information (NCBI) database with the *megablast* algorithm (McGinnis & Madden, 2004). Phylogenetic trees to investigate the clustering of individuals were built in MEGA v6 (Tamura, Stecher, Peterson, Filipski, & Kumar, 2013) to investigate the branching of pigmented individuals with known sea lamprey sequences.

### Design of environmental DNA assay: species-specific primers and probe

Due to the high taxonomic representation of mtDNA in the National Centre for Biotechnology Information (NCBI) database, we chose to design primers in conserved regions of the sea lamprey mitochondrial molecule. For that, we built a database with sequences from D-loop (control region), mtATP8 and mtATP6 genes from sea lamprey, brook lamprey and river lamprey (*Lampetra fluviatilis*). All sequences were extracted from the NCBI database. After aligning 100 lamprey sequences with the full mitogenome of sea lamprey to validate positions and confirm conservatism within the target species, primer/probe design was conducted in Primer3 (Rozen & Skaletsky, 2000).

Synthesis was outsourced to ThermoFisher Scientific^©^. After design, testing the efficiency and specificity of the primers used a three stage process: (1) confirmation of assay sensitivity in detecting various concentrations of sea lamprey DNA in TE solution; (2) confirm primer specificity in amplifying only sea lamprey DNA; and (3) verify primer efficiency to detect DNA molecules extracted from environmental samples (including in the controlled presence of sea lamprey ammocoetes).

### 1. Assay sensitivity to variable concentrations of sea lamprey DNA in solution

The efficiency and sensitivity of the designed assay was first tested against solutions of standardized DNA concentration to demonstrate that amplification occurs even under low DNA concentrations. Amplification and detection of probe fluorescence was tested at seven different DNA concentrations, obtained after 10-fold serial dilutions to the lowest concentration of 5 ng·ul^-7^ solution. Extracted DNA of adult sea lampreys was used to make dilutions (Baltazar-Soares et al. unpublished data). The assay was synthesized at ThermoFisher©, available with the ID: APWCW7W, and consists of a Taqman MGB probe (5-CACCATCTCTACTAAACAAGTT-3) labelled with the fluorescent dye FAM at the 5’-end and with a non-fluorescent quencher MGBNFQ at the 3’-end and two primers, forward: 5-GATCCTGCCCCTTGATTCTCTA-3 and reverse: 5-TCATGGTCAGGTTCAAGTGGAT-3. All quantitative real-time PCR reactions (qRT-PCR) were conducted in 20 ul reactions: 10 ul of TaqMan® Gene Expression Master Mix, 1ul assay mix and 2ul of DNA template. Thermocycler conditions were the following: holding stage at 50 °C for 2 mins, initial denaturation at 95°C for 10 mins followed by 40 cycles of denaturation at 95 °C for 15 s, and annealing at 60 °C for 1 min. All reactions were performed in triplicates, with negatives included in each plate. Note that these sequential dilutions were later used as standards in all subsequent reactions. Detection of amplified product was quantified with CT-thresholds with specialized ABI software®. All laboratorial material was exposed to UV light for 20 minutes prior to performing any protocol. The mixture of reagents and preparation of plates was done in a sterilized hood (constant-velocity) that was bleach-cleaned for 2 hours and left to dry overnight prior to work.

### 2. Validation against cross-species amplification

The occurrence of false positives compromises the credibility of eDNA tools when designed to detect specific organisms. Despite the primers and probe of the eDNA assay being designed for detecting conserved regions of the mitochondria genome, there remained a need to empirically confirm they did not also amplify DNA of other species. The control for cross-species amplification was performed on previously extracted DNA of the 16 most common freshwater fishes in the River Frome and surrounding waterbodies, with the assay applied to samples of 10 ng·ul^-1^ from brook lamprey, roach (*Rutilus rutilus*), common bream (*Abramis brama*), chub (*Squalius cephalus*), minnow (*Phoxinus phoxinus*), perch (*Perca fluviatilis*, dace (*Leuciscus leuciscus*), bleak (*Alburnus alburnus*), grayling (*Thymallus thymallus*), brown trout (*Salmo trutta*), Atlantic salmon (*Salmo salar*), gudgeon (*Gobio gobio*), European eel (*Anguilla anguilla*), carp (*Cyprinus carpio*), European barbel (*Barbus barbus*), and shad (*Alosa spp*).

### 3. Efficiency in detecting sea lamprey DNA from sediment samples

The sensitivity of the assay to detect DNA molecules in the sediment was then tested. We also inferred whether the time spent in a sample of sediment could influence efficiency of the assay. For that, we collected three ammocoetes using a dip net (250 µm mesh size), identified as sea lamprey by their pigmented caudal fin (a trait that was confirmed with molecular markers that discriminates sea lampreys ammocoetes; see Results) and placed into individual falcon tubes (50 ml). Each tube had been filled with sediment and water collected from upstream locations above in impassable barrier, where the absence of sea lamprey had been confirmed. Following the ammocoetes burying into the sediment, approximately 20 mg of sediment was removed at three-time intervals (*t*+*5, t*+*10* and *t*+*20* minutes). At the end of the process, the ammocoete was released alive to the source location. DNA was then extracted from all the collected sediment samples, alongside a negative, which comprised a sample of pure sediment prior to insertion of the lamprey. DNA extraction from the sediment samples was performed with a commercially available extraction kit (DNAeasy PowerSoil Kit©, QIAGEN) following the manufacturer’s protocol.

### 4. In-field screening of sediment deposition sites

Five field sites were targeted as monitoring locations for detecting sea lamprey ammocoetes, with the areas selected being those where sea lamprey ammocoetes were found in previous years. Sediments were collected both upstream (U) and downstream (D) of the target sediment-deposition location (C), with the objective to infer the sensitivity of the eDNA assay to detect sea lamprey DNA molecules outside the putatively optimal burrowing site. Particle size of each sediment site was estimated, in order to ensure that the sediment of C locations was composed of finer particles. Particle size was measured in a Mastersizer 3000^©^ (Malvern Panalytical) via laser light scattering of three replicates per location site.

One site per day was sampled to allow field material to be disinfected from DNA molecules with bleach overnight, with three different 4 litre trays used for each location *U, C, D* to avoid cross site contamination. Each location was sampled six times, with two of those replicates being randomly selected for DNA extraction and processing with the eDNA assay. Both the standards and each chosen sample were amplified three times. The sampling scheme with the specific geographic location of each target sediment-deposition site, plus the upstream and downstream collection points within the 13 km transect, are represented in Fig. 1.

### Statistical analyses and graphical visualization

All statistical analyses were performed in R 3.4.1 (Team, 2013). To investigate the efficiency of environmental DNA assay in relation to time in the sediment (as described in *3*), we performed a nested ANOVA with time points within specimens: *CT*_*threshold*_ *∼ specimens/time point.*

## Results

### Species identification

Sequencing of the six ammocoetes (three pigmented and three non-pigmented) produced a 419 base pair fragment and molecular inference of species-specific polymorphisms confirmed that the early stage pigmented ammocoetes were sea lamprey (Fig. 2).

**Figure 2.**
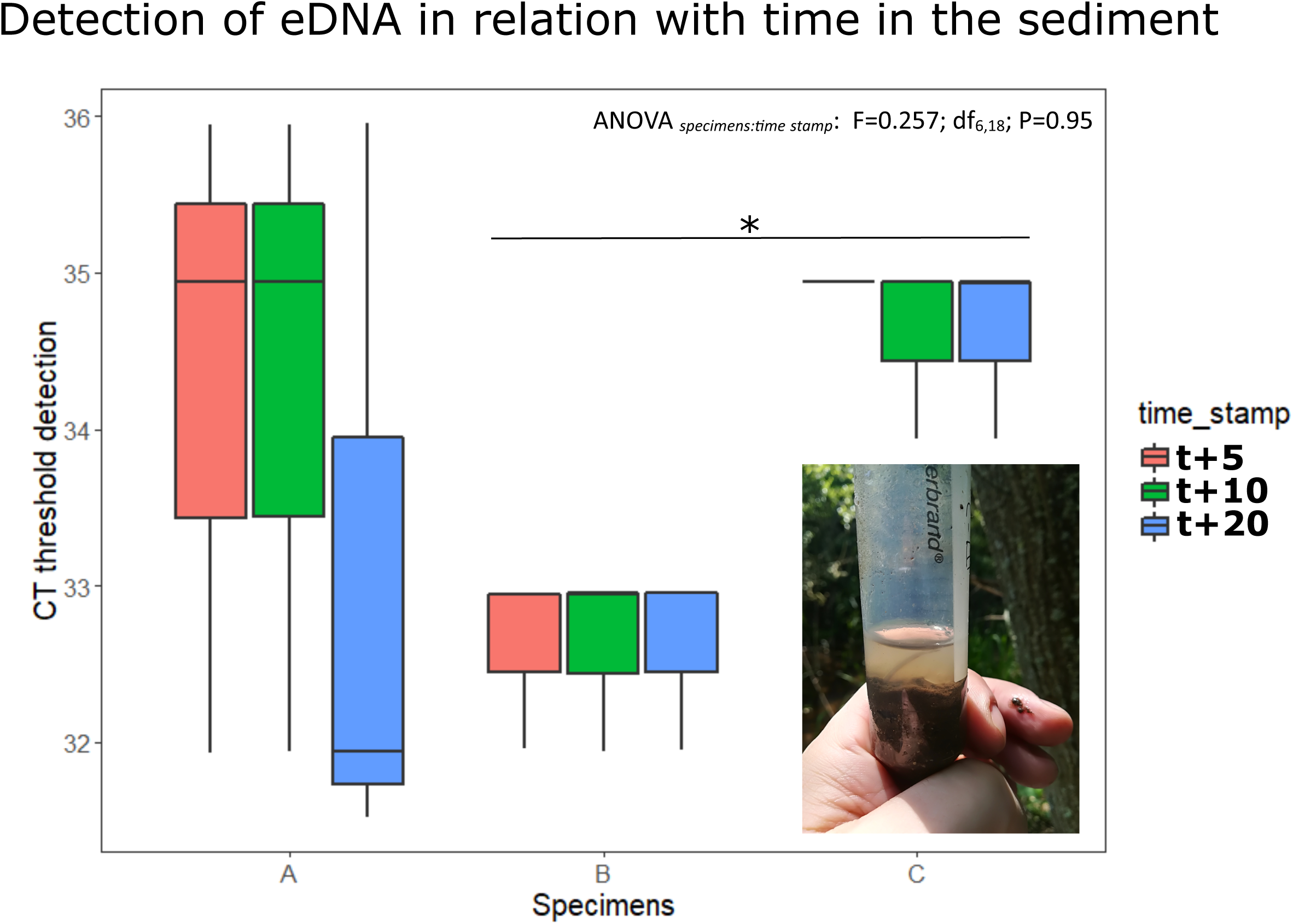
Relationship between morphological and molecular traits in the identification of sea lamprey ammocoetes. (a) Lamprey ammocoetes exhibiting the pigmentation characteristic of sea lamprey juveniles; (b) Sequences of mitochondria’s COI (mt-COI) for sea lamprey (*a* and *b*) and brook lamprey (c) as well as the picture, under microscope lens, of the respective pigmented and non-pigmented tails. Below, neighbour-joining phylogenetic tree built with mt-COI fragments.

### Development and validation of environmental DNA assay

For the 10-fold serial dilution of sea lamprey DNA, the limit of detection was 5·10 ^-5^ µg·µl^-1^, with a mean cycle threshold value (C_t_) of 37 (SD ± 0.02). The C_t_ values with DNA dilutions in later cycles (>37), corresponding to 5·10 ^-6^ µg·µl^-1^ and 5·10 ^-7^ µg·µl^-1^, were unreliable due to their probability of detection being below the 95 % confidence level. There was no cross amplification with off-target species.

### Effect of burrowing activity and time since burrowing on DNA detection

DNA molecules were detected in sediment samples after exposure with live juveniles for the three time intervals, with C_t_ ranging from 31.93 to 35.95, although there were no significant differences in terms of C_t_ detection as a response to time in the sediment (ANOVA _*specimens:time stamp*_: F_6,18_ = 0.257, P = 0.95) (Fig. 3). However, there was a significant difference between individual detection values.

**Figure 3.**
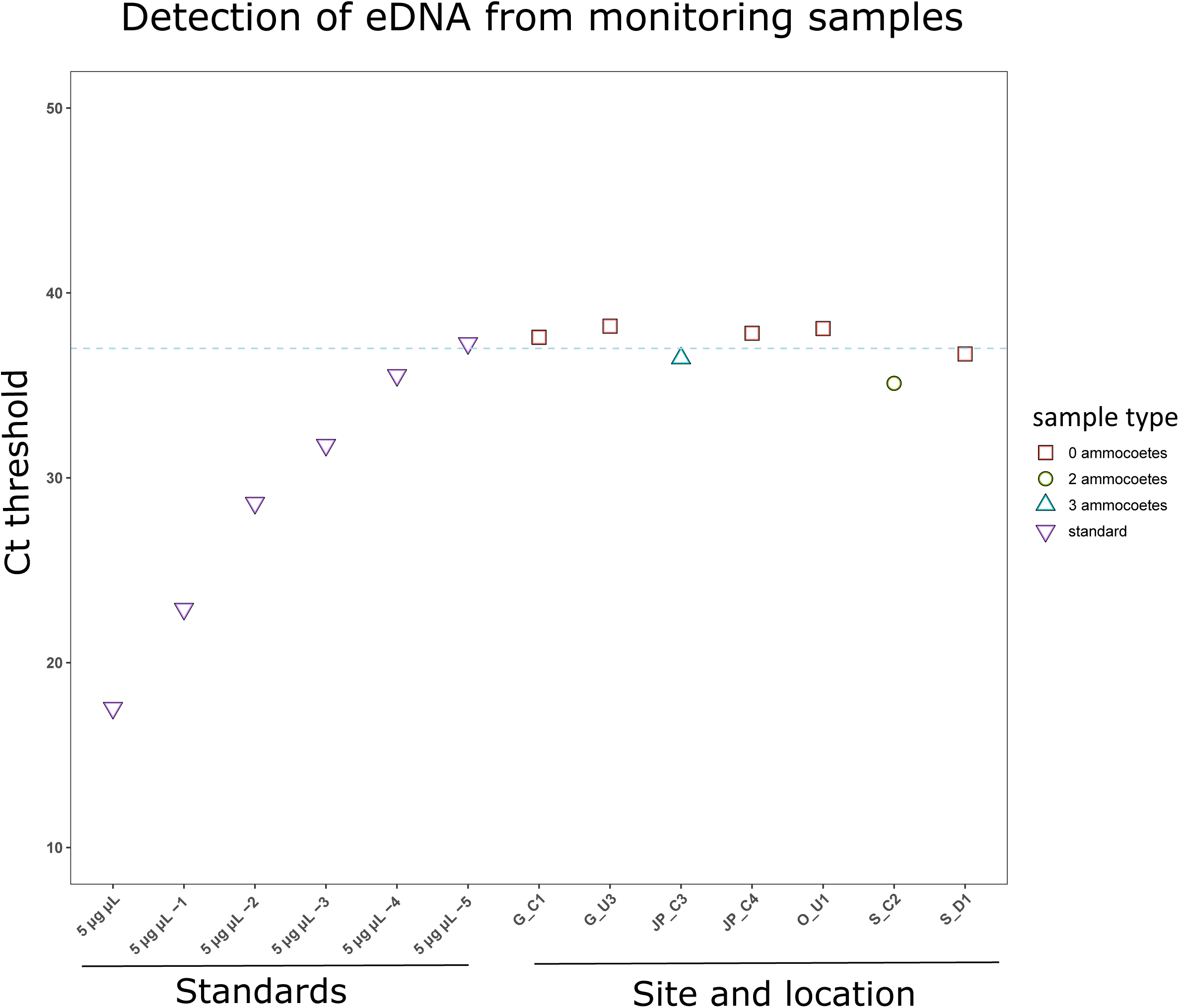
Detection of sea lamprey DNA after sediment exposure to live specimens. Variation of environmental DNA assay sensitivity to different time periods that ammocoetes where spent in the sediment. Only between individuals we found a significant difference of reported of detection threshold.

### Field application of eDNA assay to detect ammocoetes in low abundance conditions

Particles in sediment patches where sea lamprey ammocoetes were found had a mean size of 27.18µm (+/-0.68 SD), and thus smaller than those estimated from immediately upstream and downstream locations (U _*average particle size*_ = 38.72 µm, +/-1.91 SD; D _*average particle size*_ = 38.72 µm, +/-1.91 SD); ANOVA _*site:location*_: F_10,60_ = 184.68; P<0.001 (Fig S1).

The application of eDNA assay to the five river sites where ammocoetes were sampled in previous years revealed that DNA molecules of sea lamprey were detected only where ammocoetes were found on this field survey. Specifically, and despite intensive effort, we found only two ammocoetes in one of the two sampling attempts of site S (S_C2) and three ammocoetes in one of the two sampling attempts of site JP (JP_C). Application of eDNA to those samples revealed C_t_ of 35.11 and 36.46, respectively. All other samples where no ammocoetes were found during sediment sampling, as well as field negatives, had no detectable sea lamprey DNA (with C_t_ values above the detection confidence threshold; Fig. 4).

**Figure 4.**
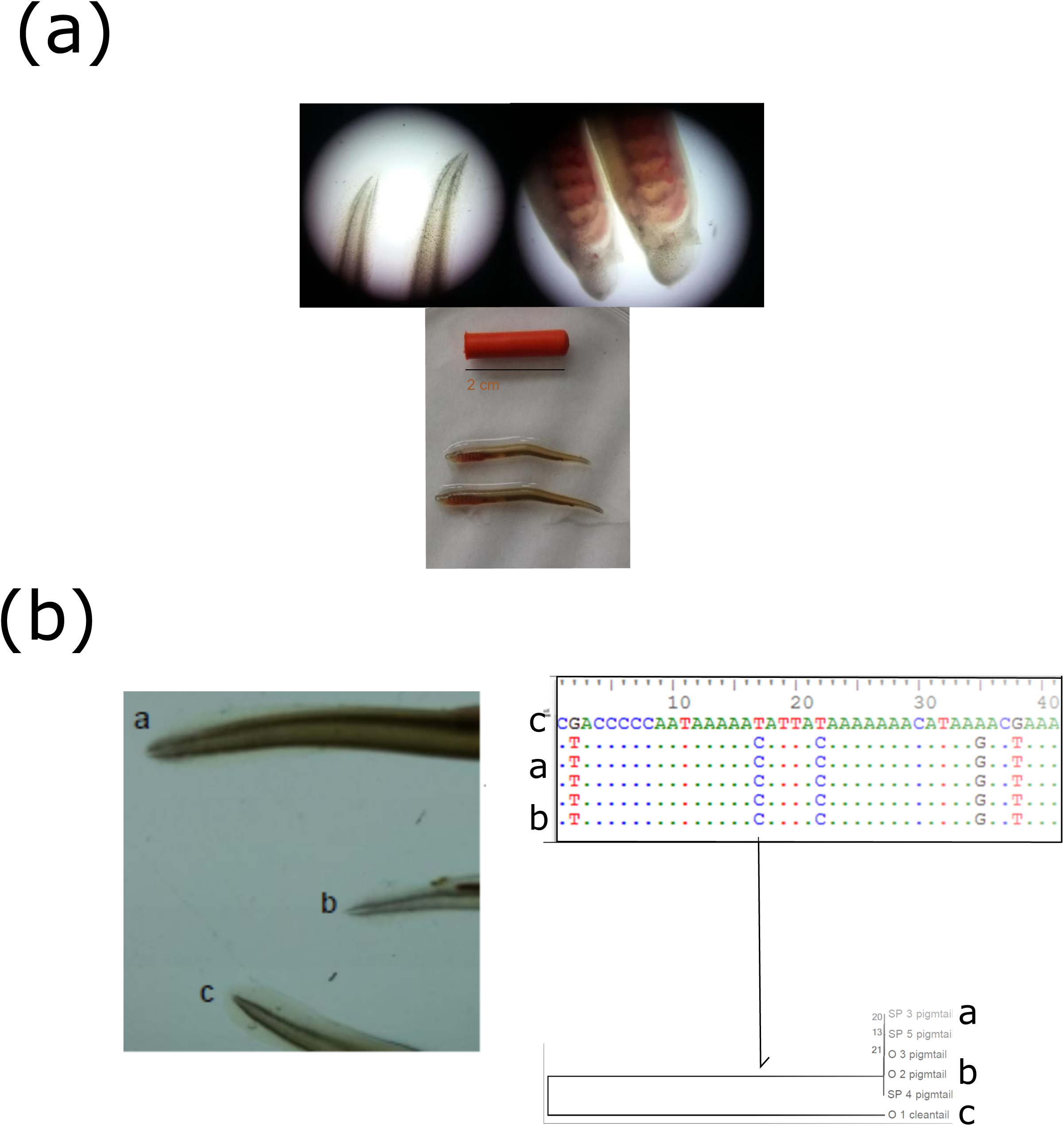
Application of environmental DNA assay to field samples. (a) Assay sensitivity to detect sea lamprey DNA in freshly collected sediment samples. Only sites where the assay amplified sea lamprey DNA are shown. Line at y(Ct)=38 depicts the standardized detection threshold.

## Discussion

Monitoring activities aiming to screen for the presence of species that utilize less conspicuous habitats is challenging. With short stays in rivers as adults to spawn but with extended development phases as larvae burrowed in sediment, monitoring of sea lamprey is challenging. Indeed, when reproduction and nesting events are rare or unable to be detected due to unsuitable river conditions, documenting the presence of this species in freshwaters must resort to trapping, dredging of sediment patches and / or electric fishing (Harvey, Noble, Nunn, Taylor, & Cowx, 2010). The advent of eDNA based detection tools came as a complement to traditional monitoring activities and, with this work, we show that ammocoete monitoring using eDNA is possible and its addition to the sea lamprey monitoring toolkit should provide considerable conservation benefits.

Our assay revealed to be highly efficient in detecting DNA molecules of sea lamprey both in standard solvents and in sediment samples. Indeed, it was able to detect the presence of sea lamprey larval DNA after a minimum exposure of only five minutes in 25 ml of sediment, with the time of exposure not inducing variation in the detection thresholds. This can be interpreted in light of their burrowing behaviours; after the initial burrowing activity, the ammocoete held still as soon as it got partially burrowed in the sediment, with stillness being characteristic of their daylight behaviour (Quintella et al., 2005). The DNA molecules detected were likely to have been shed during the intense period of burrowing activity. Whilst it has been suggested that the concentration of free DNA molecules in a given environment is a function of an individual’s metabolic activity, behaviour and abundance (Lacoursière□Roussel, Côté, Leclerc, & Bernatchez, 2016), we argue that the capacity to detect free range DNA molecules also varies with the proximity of the sampled patch to the presence of individuals. Interestingly, we further observed significant differences between individual C_t_ thresholds, suggesting that variation in individual DNA release (whether by movement, skin shedding or excretion) could also be a determining factor to consider in eDNA assays, on top of others such as density of target organisms (B. C. Stoeckle et al., 2017). Sea lamprey eDNA was detected in field samples with extremely low ammocoete abundances (with a Ct range of 35.11-36.46). This observation suggests that ammocoete density in natural sediment patches should be considered as a factor influencing detection rates of molecular material. Notably, there was no detection of qPCR reaction inhibition and all negatives were indeed negative, while positives were concomitantly tested. These results suggest that although negative results should be treated with some caution, given uncertainty around the degradation time of eDNA in the substrate (Thomsen & Willerslev, 2015b) and the duration of the ammocoete presence in sediment beds (Quintella et al., 2005), these can be overcome with further work on understanding the persistence of DNA following the movement of a species to a different location.

Recently, an eDNA assay using water samples was developed to identify key spawning areas for sea lamprey, including an investigation of the impact of physical barriers that prevent the species fully utilizing its potential spawning habitat (Bracken et al., 2019). Whilst useful for mapping the spatial distribution of spawning areas, the utility of the tool is limited by the short temporal window during which spawning occurs. Such an approach might also be inefficient when the spawning activity is lower than expected, such as in years of low adult abundance. Indeed, our sediment-based assay potentially complements the recent work of Bracken et al (2019), as it allows the screening of sediment patches downstream spawning grounds and thus estimate reproductive success.

The eDNA assay here developed will be of most benefit to ongoing sea lamprey monitoring if used as a complement to other general management practices, such as quantitative quadrat-based-or semi-quantitative electric fishing (Cowx, 2003). Furthermore, DNA-based monitoring enables increased knowledge on sea lamprey ammocoete distribution to be generated, namely related to the identification of potential nursery habitats in terms of depth. Due to methodological constrains to the application of electric fishing beyond certain depths, existing monitoring activities are only able to target sediment beds as deep as 1 m. However, recent research developments of sea lamprey ammocoete habitat utilisation suggests preferences for deeper nurseries, i.e. >2 m (Pinder et al., 2016; Taverny et al., 2012).

Through the analyses of sediment cores from deeper regions of a river, the eDNA assay also has the potential to facilitate the assessment of this species’ downstream migration to the sea and investigate the accuracy of expected successive settlement / emergence events from one sediment patch to another.

The eDNA assay here developed offers an improvement to sea lamprey current monitoring strategies. By targeting the sediment, we are proposing a holistic sea lamprey monitoring strategy able to be employed all year round to river areas that are critical for the species’ life history.

## Acknowledgments

This work was supported by a MSCA-IF (ADAPTATION) attributed to MBS. MBS is presently supported by the FCT strategic project UID/MAR/04292/2013 granted to MARE.

## Author contributions

All authors contributed to the design of the experiment, MBS, JP, WO performed the field work and completed the analyses, MBS led the writing of the manuscript, all authors contributed to revising the manuscript and all authors approved its submission.

## Bibliography

Almada, V. C., Pereira, A. M., Robalo, J. I., Fonseca, J. P., Levy, A., Maia, C., & Valente, A. (2008). Mitochondrial DNA fails to reveal genetic structure in sea-lampreys along European shores. Molecular Phylogenetics and Evolution, 391–396.

Antognazza, C. M., Britton, J. R., Potter, C., Franklin, E., Hardouin, E. A., Gutmann Roberts, C., Andreou, D. (2019). Environmental DNA as a non-invasive sampling tool to detect the spawning distribution of European anadromous shads (Alosa spp.). Aquatic Conservation: Marine and Freshwater Ecosystems, 29(1), 148–152.

Bohmann, K., Evans, A., Gilbert, M. T. P., Carvalho, G. R., Creer, S., Knapp, M., De Bruyn, M. (2014). Environmental DNA for wildlife biology and biodiversity monitoring. Trends in Ecology & Evolution, 29(6), 358–367.

Bracken, F. S., Rooney, S. M., Kelly-Quinn, M., King, J. J., & Carlsson, J. (2019). Identifying spawning sites and other critical habitat in lotic systems using eDNA “snapshots”: A case study using the sea lamprey Petromyzon marinus L. Ecology and Evolution, 9(1), 553–567.

Cowx, J. H. I. (2003). Monitoring the River, Brook and Sea Lamprey, Lampetra fluviatilis, L. planeri and Petromyzon marinus. Retrieved from Peterborough:

Dias, M. S., Tedesco, P. A., Hugueny, B., Jézéquel, C., Beauchard, O., Brosse, S., & Oberdorff, T. (2017). Anthropogenic stressors and riverine fish extinctions. Ecological indicators, 79, 37–46.

Dodds, W. K., Perkin, J. S., & Gerken, J. E. (2013). Human impact on freshwater ecosystem services: a global perspective. Environmental science & technology, 47(16), 9061–9068.

Doi, H., Inui, R., Akamatsu, Y., Kanno, K., Yamanaka, H., Takahara, T., & Minamoto, T. (2017). Environmental DNA analysis for estimating the abundance and biomass of stream fish. Freshwater Biology, 62(1), 30–39.

Evans, N. T., Shirey, P. D., Wieringa, J. G., Mahon, A. R., & Lamberti, G. A. (2017). Comparative cost and effort of fish distribution detection via environmental DNA analysis and electrofishing. Fisheries, 42(2), 90–99.

Ficetola, G. F., Miaud, C., Pompanon, F., & Taberlet, P. (2008). Species detection using environmental DNA from water samples. Biology Letters, 4(4), 423–425.

Guo, Z., Andreou, D., & Britton, J. (2017). Sea lamprey Petromyzon marinus biology and management across their native and invasive ranges: promoting conservation by knowledge transfer. Reviews in Fisheries Science & Aquaculture, 25(1), 84–99.

Hall, T. A. (1999). BioEdit: a user-friendly biological sequence alignment editor and analysis program for Windows 95/98/NT. Paper presented at the Nucleic acids symposium series.

Harvey, J. P., Noble, R. A., Nunn, A. D., Taylor, R. J., & Cowx, I. G. (2010). Monitoring sea lamprey Petromyzon marinus ammocoetes in SAC rivers: a case study on the River Wye. In Conservation Monitoring in Freshwater Habitats (pp. 193–206): Springer.

Kelly, F. L., & King, J. J. (2001). A review of the ecology and distribution of three lamprey species, Lampetra fluviatilis (L.), Lampetra planeri (Bloch) and Petromyzon marinus (L.): a context for conservation and biodiversity considerations in Ireland. Paper presented at the Biology and Environment: Proceedings of the Royal Irish Academy.

Lacoursière-Roussel, A., Côté, G., Leclerc, V., & Bernatchez, L. (2016). Quantifying relative fish abundance with eDNA: a promising tool for fisheries management. Journal of Applied Ecology, 53(4), 1148–1157.

Mateus, C. S., Rodríguez-Muñoz, R., Quintella, B. R., Alves, M. J., & Almeida, P. R. (2012). Lampreys of the Iberian Peninsula: distribution, population status and conservation. Endangered Species Research, 16(2), 183–198.

McGinnis, S., & Madden, T. L. (2004). BLAST: at the core of a powerful and diverse set of sequence analysis tools. Nucleic Acids Research, 32(suppl_2), W20–W25.

Pinder, A. C., Hopkins, E., Scott, L. J., & Britton, J. R. (2016). Rapid visual assessment of spawning activity and associated habitat utilisation of sea lamprey (Petromyzon marinus Linnaeus, 1758) in a chalk stream: implications for conservation monitoring. Journal of Applied Ichthyology, 32(2), 364–368.

Quintella, B., Andrade, N., Espanhol, R., & Almeida, P. (2005). The use of PIT telemetry to study movements of ammocoetes and metamorphosing sea lampreys in river beds. Journal of Fish Biology, 66(1), 97–106.

Radinger, J., Britton, J. R., Carlson, S. M., Magurran, A. E., Alcaraz-Hernández, J. D., Almodóvar, A., Oliva-Paterna, F. J. (2019). Effective monitoring of freshwater fish. Fish and Fisheries.

Rodríguez-Muñoz, R., Waldman, J., Grunwald, C., Roy, N., & Wirgin, I. (2004). Absence of shared mitochondrial DNA haplotypes between sea lamprey from North American and Spanish rivers. Journal of Fish Biology, 64(3), 783–787.

Rozen, S., & Skaletsky, H. (2000). Primer3 on the WWW for general users and for biologist programmers. In Bioinformatics methods and protocols (pp. 365–386): Springer.

Sorensen, P. W., & Vrieze, L. A. (2003). The chemical ecology and potential application of the sea lamprey migratory pheromone. Journal of Great Lakes Research, 29, 66–84.

Stoeckle, B. C., Beggel, S., Cerwenka, A. F., Motivans, E., Kuehn, R., & Geist, J. (2017). A systematic approach to evaluate the influence of environmental conditions on eDNA detection success in aquatic ecosystems. PLoS One, 12(12), e0189119–e0189119. doi:10.1371/journal.pone.0189119

Stoeckle, M. Y., Soboleva, L., & Charlop-Powers, Z. (2017). Aquatic environmental DNA detects seasonal fish abundance and habitat preference in an urban estuary. PLoS One, 12(4), e0175186.

Tamura, K., Stecher, G., Peterson, D., Filipski, A., & Kumar, S. (2013). MEGA6: molecular evolutionary genetics analysis version 6.0. Molecular Biology and Evolution, 30(12), 2725–2729.

Taverny, C., Lassalle, G., Ortusi, I., Roqueplo, C., Lepage, M., & Lambert, P. (2012). From shallow to deep waters: habitats used by larval lampreys (genus Petromyzon and Lampetra) over a western European basin. Ecology of Freshwater Fish, 21(1), 87–99.

Team, R. C. (2013). R: A language and environment for statistical computing.

Thomsen, P. F., Kielgast, J., Iversen, L. L., Wiuf, C., Rasmussen, M., Gilbert, M. T. P., Willerslev, E. (2012). Monitoring endangered freshwater biodiversity using environmental DNA. Molecular Ecology, 21(11), 2565–2573.

Thomsen, P. F., & Willerslev, E. (2015a). Environmental DNA–An emerging tool in conservation for monitoring past and present biodiversity. Biological Conservation, 183, 4–18.

Thomsen, P. F., & Willerslev, E. (2015b). Environmental DNA – An emerging tool in conservation for monitoring past and present biodiversity. Biological Conservation, 183, 4–18. doi: https://doi.org/10.1016/j.biocon.2014.11.019

