## Supplementary figures and images for "A non-invasive eDNA tool for detecting sea lamprey larvae in river sediments: analytical validation and field testing in a low abundance ecosystem"

### Fig.S1

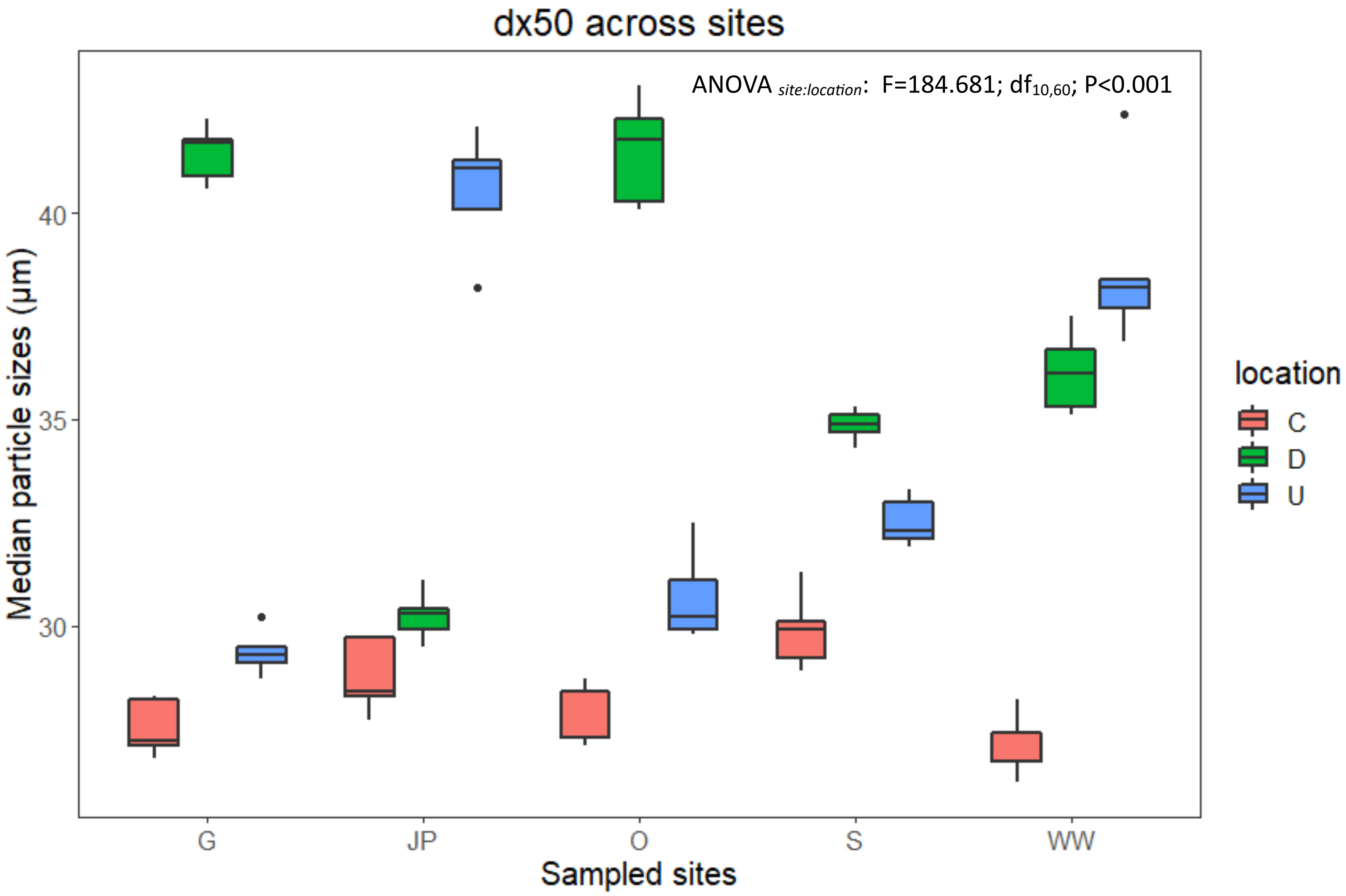
